# Genetic structure is stronger across human-impacted habitats than among islands in the coral *Porites lobata*

**DOI:** 10.1101/574665

**Authors:** Kaho H. Tisthammer, Zac H. Forsman, Robert J. Toonen, Robert H. Richmond

**Affiliations:** Kewalo Marine Laboratory, University of Hawai‘i at Mānoa, Honolulu, Hawai‘i, United States of America; Department of Biology, San Francisco State University, United States of America; Hawaiʻi Institute of Marine Biology, University of Hawai‘i at Mānoa, Kāneʻohe, Hawai‘i, United States of America

## Abstract

We examined genetic structure in the lobe coral *Porites lobata* among pairs of highly variable and high-stress nearshore sites and adjacent less variable and less impacted offshore sites on the islands of Oʻahu and Maui, Hawai‘i. Using an analysis of molecular variance framework, we tested whether populations were more structured by geographic distance or environmental extremes. The genetic patterns we observed followed isolation by environment, where nearshore and adjacent offshore populations showed significant genetic structure at both locations (AMOVA *F_ST_* = 0.04 ∼ 0.19, *P* < 0.001), but no significant isolation by distance between islands. In contrast, a third site with a less impacted nearshore site showed no significant structure. Strikingly, corals from the two impacted nearshore sites on different islands over 100km apart with similar environmentally stressful conditions were genetically closer (*F_ST_*∼ 0, P = 0.733) than those within a single location less than 2 km apart (*F_ST_*= 0.041∼0.079, P < 0.01). Our results suggest that ecological boundaries appear to play a strong role in forming genetic structure in the coastal environment, and that genetic divergence in the absence of geographical barriers to gene flow may be explained by disruptive selection across contrasting habitats.

## Introduction

Coral reefs are centers of marine biodiversity and productivity that provide a variety of ecosystem services of substantial cultural and economic value to humankind, yet coral reefs worldwide are under serious threat as a result of human activities^1,2^. Average global coral cover has declined dramatically in the past 100 years due to a range of impacts such as sedimentation, pollution, overfishing, disease outbreaks and climate change^1–3^.

Such effects are particularly pronounced in nearshore marine habitats, which are increasingly exposed to reduced water quality due to human activities^4^. Recent rapid coastal development, along with coastal industrial and recreational activities, have resulted in introducing sediments, nutrients and a variety of chemical pollutants to the nearshore environments^4–6^. These local stressors often create a steep environmental gradient of water quality from nearshore toward offshore areas, and ‘signs of coral health impairment’ are usually detected along with the gradient (e.g.^5,7,8^). Additionally, very nearshore marine habitats are naturally exposed to higher fluctuations in temperature, pH and other environmental variables, creating contrasting environmental conditions relative to more stable offshore environments^9,10^. Some corals, however, continue to thrive in such nearshore ‘suboptimal’ habitat^11^, indicating that these individuals can withstand such stressors.

What impact does an ecological landscape with such a strong gradient have on the genetics of the organisms? ‘Isolation by environment’ (IBE^12^) describes a pattern in which genetic differentiation increases with environmental differences, independent of geographic distance. Isolation by environment is a process that emphasizes the role of environmental heterogeneity and ecology in forming genetic structure, likely because of natural selection, in contrast to ‘isolation by distance’ (IBD^13^), which predicts that the degree of genetic differentiation increases with geographic distance due primarily to dispersal limits^14^. Importantly, IBD is a neutral process in which dispersal limits gene flow and the scale over which genetic structure accumulates, whereas IBE explicitly takes into account environmental differences among sites^12,15^. Isolation by environment can be generated by different processes, including natural selection, sexual selection, reduced hybrid fitness, and biased dispersal; examples of the terms describing a specific case of IBE include ‘isolation by adaptation’ (IBA^16^), ‘isolation by colonization’ (IBC^17^), and ‘isolation by resistance’ (IBR^18^). IBA and IBC emphasize the role of selection in forming genetic structure, and IBR describes correlation of genetic distance and resistance distance (i.e. friction to dispersal)^18^. IBE along with related terms result in a pattern where genetic distance increases as ecological distance increases, but not with geographic distance for most loci^12,15^. Theoretically, IBA, IBC and other processes will result in different distributions of genetic variation across landscapes^15^, though in reality, multiple processes almost always contribute to structuring genetic variation, and pinpointing the possible underlying processes may be difficult^14^. For coastal marine ecosystems, often the distances between the impacted nearshore and un-impacted offshore sites are relatively small with no apparent dispersal barrier between adjacent sites for broadcast spawning species with pelagic larval development, providing an excellent opportunity to study IBE.

Carlon and Budd ^19^ described a pair of incipient species in the coral *Favia fragum* associated with strong ecological gradients. The two types are largely restricted to alternate seagrass and adjacent coral reef habitats, but retain phenotypic distinction in a narrow zone of ecological overlap ^19^. Subsequent work showed that the morphologies were heritable, and selection appeared to limit gene flow between the ectomorphs^20^. Carlon *et al*.^20^ postulated that divergent selection for “Tall” and “Short” ectomorphs of these inbreed and brooding corals was driving the diversification of this coral via an ecological model of speciation (*sensu*^21^). Genetic divergence across nearshore and offshore habitats has also been observed for broadcast spawning corals; for example in *Seriatopora hystrix* in Australia^22^, and *Porites lobata* in American Samoa^23^. Here we pose the question of whether there is reason to believe such a pattern might be observed in contrasting habitats in Hawai‘i, to add to a growing number of studies indicating that similar patterns may be more ubiquitous than previously assumed.

Maunalua Bay, Hawai‘i, O‘ahu, was selected as a study site due to the existence of a strong environmental gradients; large-scale urbanization in adjacent watersheds has caused severe deterioration in the health and extent of its nearshore coral reefs over the last century^24^. Corals that survive in these affected nearshore areas are under chronic stress, and a previous survey showed significantly different cellular stress responses of individual colonies along the environmental gradient of pollutants and sedimentation from the inner bay toward offshore^24–26^ (Fig S1). *Porites lobata* (Dana, 1846), the study species, occurs over a wide geographic range in the tropical Pacific Ocean^27^, and several studies have documented a pattern of isolation by distance across archipelagic^28^ or broader scales ^29^, with little evidence of restricted gene flow among geographically proximate reefs at inter-island distances (but see^23^). This massive coral is also known for its robustness; for example, *P. lobata* shows a high tolerance for sedimentation^30^ and bleaching^31^, and a colony can recover from partial mortality due to tissues residing deep within the perforate skeleton, a phenomenon referred to as the ‘Phoenix effect’^32^. *Porites lobata* is one of the most dominant scleractinian coral species in Hawai‘i^33^. Additionally, *P. lobata* shows high fidelity to a specific endo-symbiont, *Symbiodinium* Clade C15^23, 34–36^, which allows us to focus on responses of the host coral to environmental differences.

At the nearshore site of Maunalua Bay, the suspended sediment concentration periodically exceeds several hundred mg/L, and the run-off water introduces toxicants such as benzo[a]pyrene, benzo[k]fluoranthene, phenanthrene and alpha-chlordane (Fig. 1, Fig. S1)^24,37,38^. The temperature, salinity and turbidity likewise all show higher fluctuations and gradients across the bay^38^.The distance between the studied offshore and nearshore sites were less than 2 km across the extent of this gradient, and water movement in the areas suggests no dispersal barrier between the sites^33^.

**Fig. 1.**
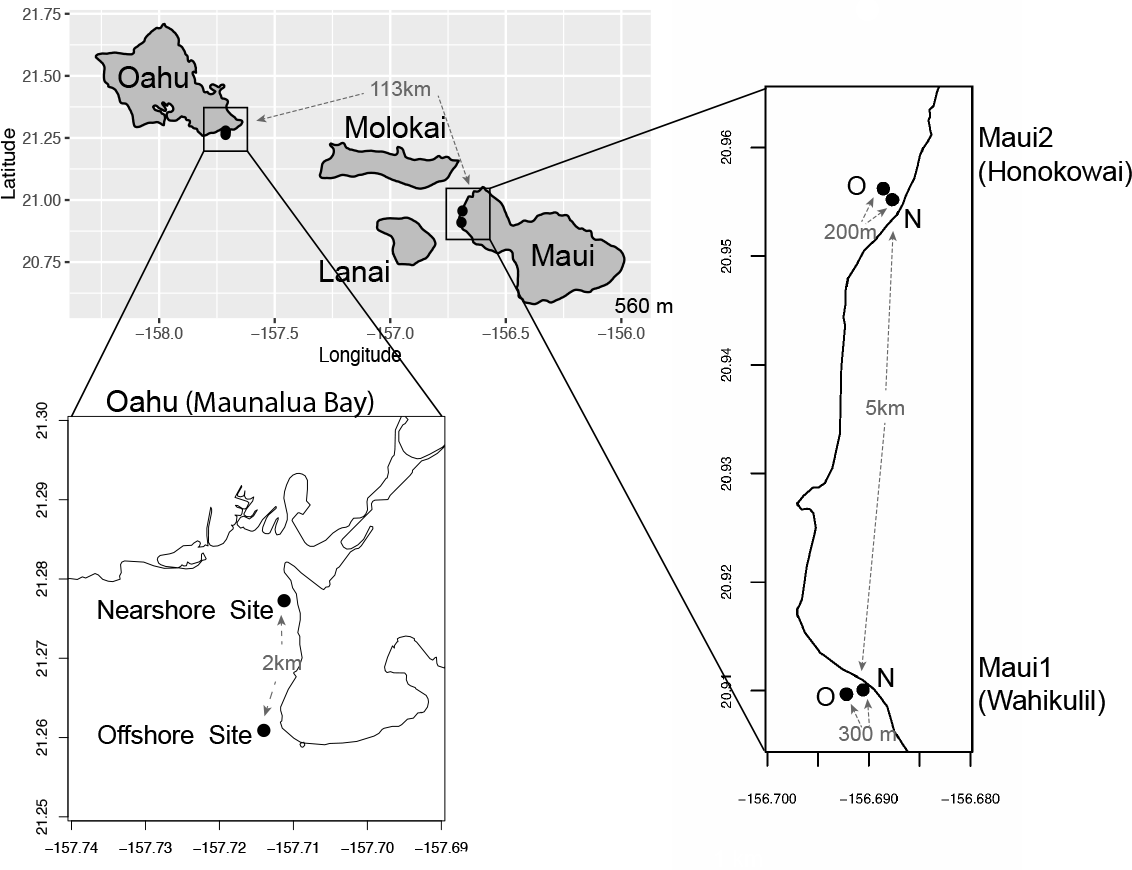
Maps of sampling locations. The distances between the sampling sites are shown in gray.

Similarly, the coral reefs off West Maui have experienced a dramatic decline in their coral cover from land-based anthropogenic impacts over the last several decades^39^. Substantial deterioration in the health of West Maui’s coral reefs has lead Wahikuli and Honokōwai watersheds of West Maui to be designated as priority sites for conservation and management by the United States Coral Reef Task Force (USCRTF) and the State of Hawai‘i^40^. The Wahikuli study site is directly exposed to terrestrial run-off, due to its topography and current patterns ^41^, causing high turbidity especially after heavy rains. Despite their proximity, the nearshore area at Wahikuli has markedly different water quality than offshore reefs roughly 300m away (Fig. S2). In contrast, the nearshore area at the Honokōwai site is less affected by runoff, because it does not receive any direct stream discharge, resulting in consistently lower turbidity than the Wahikuli nearshore site^42^.

Differences in water quality and sedimentation loads of nearshore and offshore environments in Maunalua Bay, Oʻahu and Wahikuli, Mauʻi represent strong gradients of anthropogenic impacts that create highly contrasting environments in close proximity. Therefore, we undertook a genetic analysis of *P. lobata* across these sites to explore the possibility of isolation by environment. By comparing corals collected from heavily impacted nearshore environments to nearby congeners from more oceanic conditions, we sought to distinguish the roles played by ecology and anthropogenic impacts to the environment on the genetics of coral populations, in contrast to geographical distance limiting dispersal among similar habitats on adjacent islands. We predicted that the genetic structure of coral populations from areas with a strong anthropogenic impact gradient would follow IBE, rather than IBD. *Porites lobata* populations from Maunalua Bay, Oʻahu (hereafter Oʻahu) and Wahikuli, Maui (hereafter Maui1) represented the sites with such a strong environmental gradient, while Honokōwai, Maui (hereafter Maui2) represents a similar nearshore-offshore comparison site, but without a strong anthropogenic gradient (Fig. 1). At each site, we assessed the degrees of genetic differentiation and genetic diversity of *P. lobata* between adjacent strongly anthropogenically impacted ‘high-stress’ nearshore and ‘low-stress’ offshore sites, and compared them within and between locations to understand the effects of habitat types, anthropogenic impacts, and geographical distance on the genetic structure of reef building corals.

## Results

### Nearshore vs. offshore comparison of genetic structure and diversity *of P. lobata* populations

#### Oʻahu (Maunalua Bay)

For Oʻahu *P. lobata* populations, the degree of genetic differentiation was estimated using analysis of molecular variance (AMOVA^43^) between the nearshore and offshore sites using three genetic markers; existing nuclear ITS1-5.8S-ITS2 region (ITS), mitochondrial putative control region (CR), and novel nuclear histone region spanning H2A to H4 (H2), developed for this study. The AMOVA results for both nuclear makers revealed clear genetic differentiation between the two sites (ITS, *F_ST_* = 0.1918, *P* < 0.001; H2, *F_ST_*= 0.0715, *P* < 0.001) (Table 1). The mitochondrial marker (CR) did not detect significant differentiation (*F_ST_* = 0.086, *P* = 0.148), which was not surprising due to its extremely low variability in corals and cnidarians in general 44. The numbers of shared haplotypes (alleles) between the nearshore and offshore Oʻahu populations were also low; out of 37 ITS haplotypes identified from the 70 total sequences, only three (8.1%) were shared between the sites. For H2, there were 54 unique haplotypes out of 86 total phased sequences, and only 5 sequences (9.3%) were shared between the sites (Table 2). The pattern of genetic structure was visualized using network analysis, which revealed sequences clustering into three major groups in both ITS and H2 markers, which consisted of one cluster dominated by the nearshore individuals, the second one dominated by the offshore individuals, and the last group with approximately mixed origins (Fig. 2). For CR, three haplotypes were identified from 27 sequences, all of which were present at both sites. Interestingly, the most common haplotype was the most dominant one at the nearshore site, while the second common haplotype was the dominant haplotype at the offshore site, though the AMOVA results were not significant (Fig. 3).

**Fig. 2.**
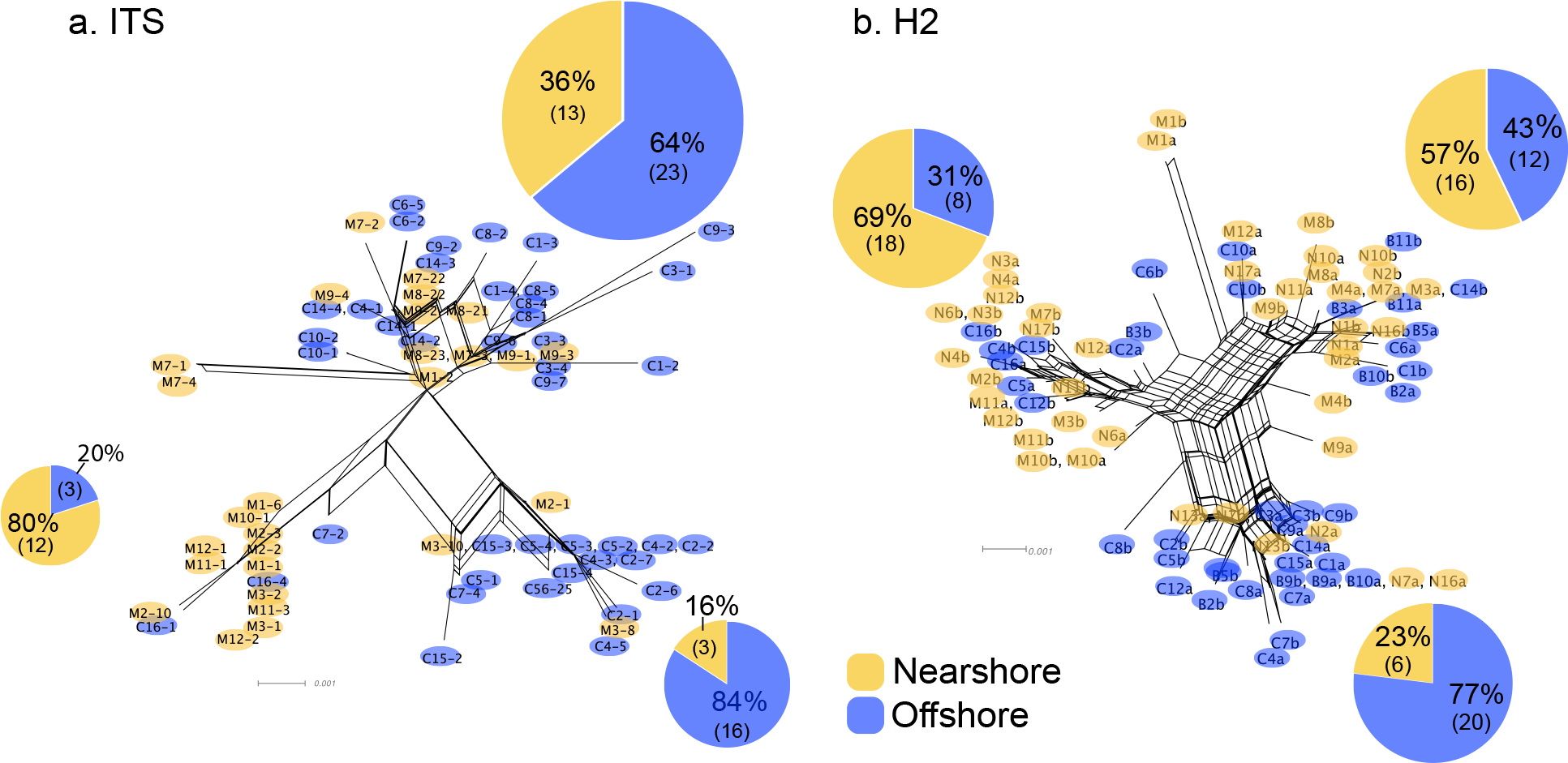
Diagrams of neighbor-net tree networks generated by SplitsTree v.4.14.2 for O‘ahu (Maunalua Bay) *P. lobata* populations, based on (a) ITS and (b) H2. Pie charts represent the proportion of sequences in each cluster.

**Table 1.** AMOVA results of *P. lobata* from Oʻahu (Maunalua Bay).

| | Source of Variation | Variance components | % Variance | $F_{ST}$ |
| --- | --- | --- | --- | --- |
| ITS<br>(n=70) | Between populations | 2.27 | 19.18 | **0.1918\*\*\*** |
|  | Within populations | 9.56 | 80.82 |  |
| H2<br>(n=43) | Between populations | 0.29 | 7.15 | **0.0715\*\*\*** |
|  | Within populations | 1.30 | 31.88 |  |
|  | Within individuals | 2.49 | 60.96 |  |
| CR<br>(n=20) | Between populations | 0.034 | 8.49 | 0.08595<br>($P$ = 0.148) |
|  | Within populations | 0.370 | 91.5 |  |

**Table 2.**
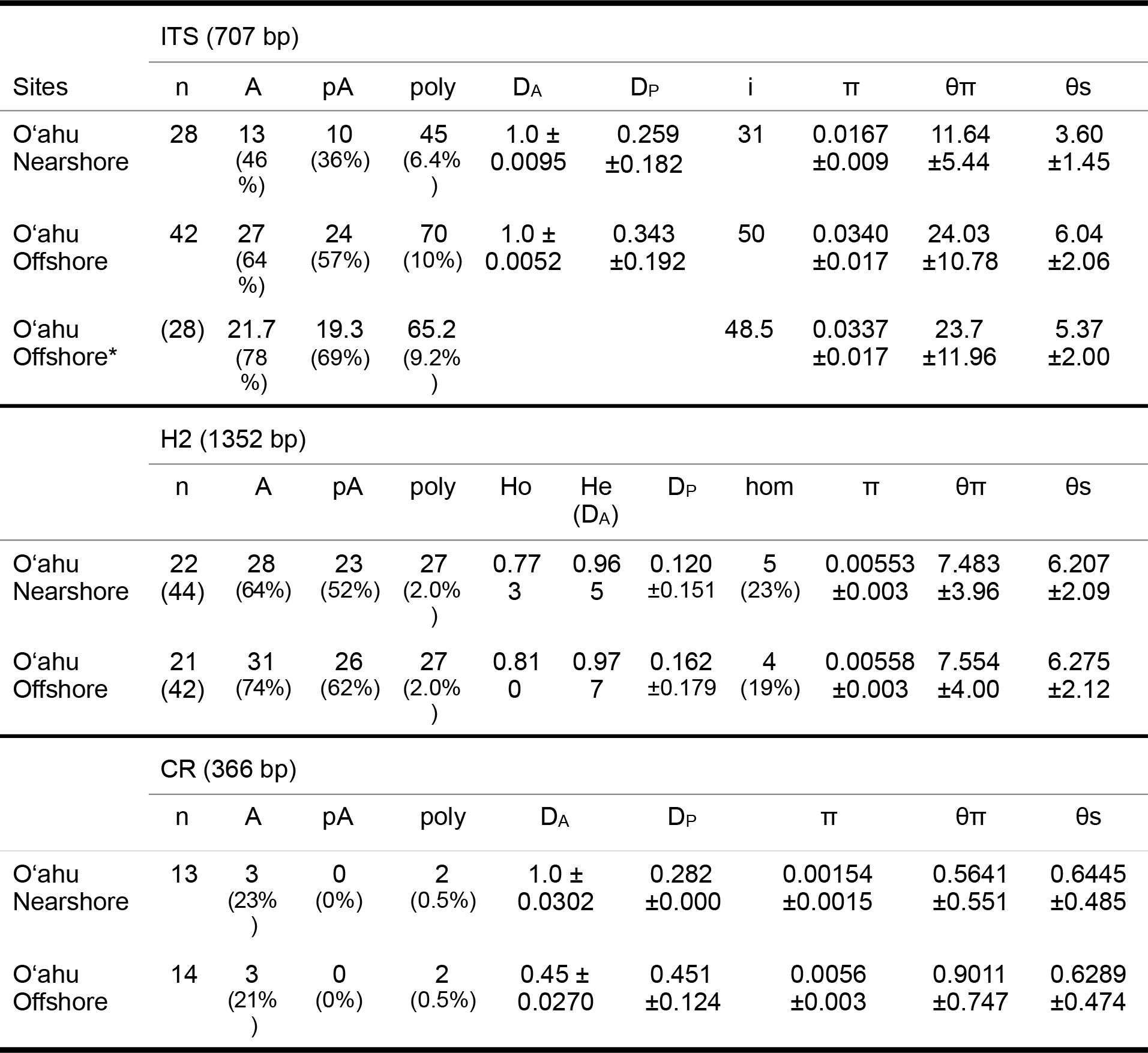
Population genetic statistics of *P. lobata* from Oʻahu (Maunalua Bay). Sample size (n, for H2, the number in () represents the number of phased sequences.), number of haplotypes (A), number of private haplotypes (pA), number of polymorphic sites (poly), mean overall gene diversity (D_A_ ± SD), mean gene diversity for polymorphic sites only (D_P_ ± SD), observed heterozygosity (H_o_), expected heterozygosity (He), number of indels (i), number of homozygous individuals (hom), nucleotide diversity (π ± SD), theta estimator 1 (θπ: expected heterozygosity at a nucleotide position estimated from the mean π), theta estimator 2 (Watterson estimator, θs). *Standardized values to the minimum sample size of 28.

| ITS (707 bp) |  |  |  |  |  |  |  |  |  |  |  |
| --- | --- | --- | --- | --- | --- | --- | --- | --- | --- | --- | --- |
| Sites | n | A | pA | poly | D<sub>A</sub> | D<sub>P</sub> | i | π | θπ | θs |  |
| O'ahu Nearshore | 28 | 13<br>(46%) | 10<br>(36%) | 45<br>(6.4%) | 1.0 ±<br>0.0095 | 0.259<br>±0.182 | 31 | 0.0167<br>±0.009 | 11.64<br>±5.44 | 3.60<br>±1.45 |  |
| O'ahu Offshore | 42 | 27<br>(64%) | 24<br>(57%) | 70<br>(10%) | 1.0 ±<br>0.0052 | 0.343<br>±0.192 | 50 | 0.0340<br>±0.017 | 24.03<br>±10.78 | 6.04<br>±2.06 |  |
| O'ahu Offshore* | (28) | 21.7<br>(78%) | 19.3<br>(69%) | 65.2<br>(9.2%) |  |  | 48.5 | 0.0337<br>±0.017 | 23.7<br>±11.96 | 5.37<br>±2.00 |  |
| H2 (1352 bp) |  |  |  |  |  |  |  |  |  |  |  |
|  | n | A | pA | poly | Ho | He<br>(D<sub>A</sub>) | D<sub>P</sub> | hom | π | θπ | θs |
| O'ahu Nearshore | 22<br>(44) | 28<br>(64%) | 23<br>(52%) | 27<br>(2.0%) | 0.77<br>3 | 0.96<br>5 | 0.120<br>±0.151 | 5<br>(23%) | 0.00553<br>±0.003 | 7.483<br>±3.96 | 6.207<br>±2.09 |
| O'ahu Offshore | 21<br>(42) | 31<br>(74%) | 26<br>(62%) | 27<br>(2.0%) | 0.81<br>0 | 0.97<br>7 | 0.162<br>±0.179 | 4<br>(19%) | 0.00558<br>±0.003 | 7.554<br>±4.00 | 6.275<br>±2.12 |
| CR (366 bp) |  |  |  |  |  |  |  |  |  |  |  |
|  | n | A | pA | poly | D<sub>A</sub> | D<sub>P</sub> | π | θπ | θs |  |  |
| O'ahu Nearshore | 13 | 3<br>(23%) | 0<br>(0%) | 2<br>(0.5%) | 1.0 ±<br>0.0302 | 0.282<br>±0.000 | 0.00154<br>±0.0015 | 0.5641<br>±0.551 | 0.6445<br>±0.485 |  |  |
| O'ahu Offshore | 14 | 3<br>(21%) | 0<br>(0%) | 2<br>(0.5%) | 0.45 ±<br>0.0270 | 0.451<br>±0.124 | 0.0056<br>±0.003 | 0.9011<br>±0.747 | 0.6289<br>±0.474 |  |  |

**Fig. 3.**
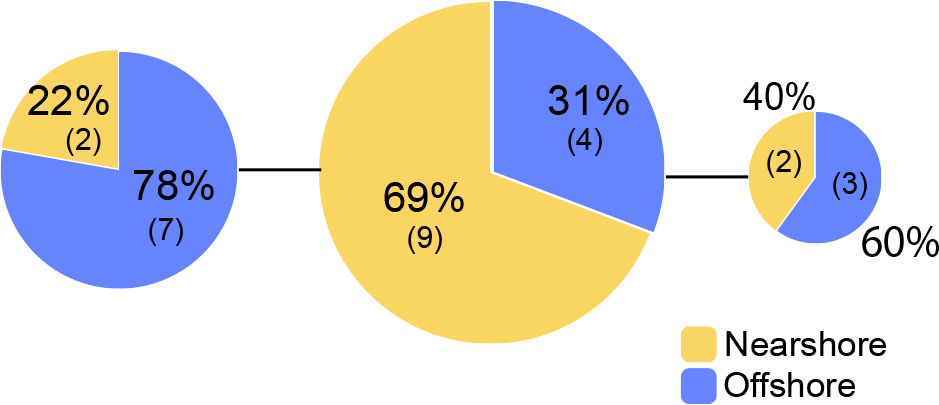
Haplotype network using the mitochondrial putative control region (CR) for the O‘ahu (Maunalua Bay) *P. lobata* populations.

The pattern of genetic diversity also differed between the nearshore and offshore populations. The degree of genetic diversity was higher at the offshore site; percent private alleles (pA), percent polymorphic sites (poly), and nucleotide diversity level (π) were almost twice as high in the offshore population as in the nearshore one based on ITS (Table 2). Standardizing sample size by random resampling confirmed that this was not an artifact of a larger sample size of the offshore population (Table 2). Rarefaction analysis of ITS sequences also confirmed that allelic richness of the offshore population (Richness = 21.6 ± 2.1 at n=28, 95% CI, 18.8 to 24.5) was clearly higher than that of the nearshore population (Richness = 13) (Fig.4). The level of genetic diversity in H2 was also higher in the offshore populations, but the difference was not as large as in ITS; the number of haplotypes (A), the number of private allele (pA), the heterozygosity level (H_O_), the number of heterozygous individuals, and mean gene diversity (D_A,_ D_P_) all had marginally higher values in the offshore samples (Table 2). In both nuclear markers, θπ (the expected heterozygosity estimated from the average nucleotide diversity) was higher than θs (the theta estimated from the number of segregating sites).

**Fig. 4.**
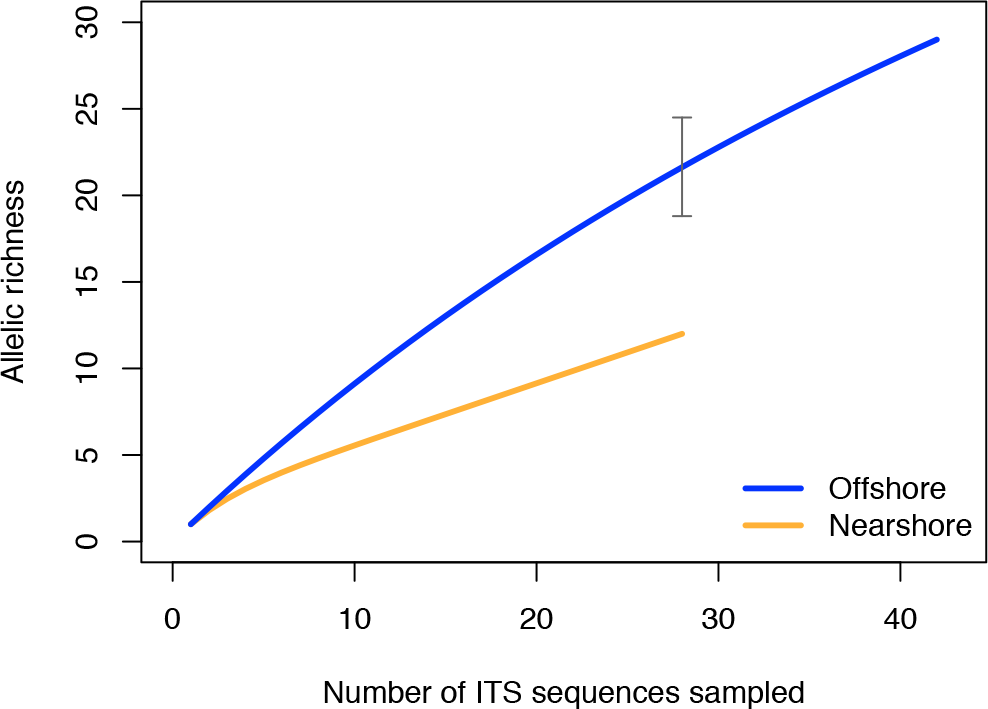
Rarefaction curve of allelic richness of ITS sequences from the *P. lobata* nearshore and offshore populations of Maunalua Bay, Hawaiʻi. The gray bar indicates the 95% confidence interval of the allelic richness estimation for the offshore population at n=28.

#### Maui

At the two study locations on the island of Maui, patterns of genetic structure of *P. lobata* populations between the nearshore and offshores sites were analyzed using the novel H2 marker (see Material and Methods). At Maui1, where a strong environmental gradient exists, significant genetic differentiation was detected (*F_ST_* = 0.0415, *P* = 0.0308), but the level of genetic diversity was comparable between the two sites (Table 3). At Maui2, which had much less contrasting environmental conditions between the nearshore and offshore sites, the AMOVA results found no significant genetic differentiation between the nearshore and offshore sites (*F_ST_* = 0.0019, *P* = 0.991). The level of genetic diversity at Maui2 appeared slightly higher in the nearshore population, which had higher numbers of haplotypes (A), polymorphic sits (poly), and heterozygous individuals (Table 3). The theta estimators of Maui2 also showed a different pattern from the Oʻahu and Maui1 populations, with higher values of θs than those of θπ at both nearshore and offshore sites.

**Table 3.** Population genetic statistics of ***P. lobata*** from Maui1 and Maui2 sites based on H2 (1319 bp). See Table 2 for symbols and abbreviations.

|  | H2 (1221 bp) |  |  |  |  |  |  |  |  |  |
| --- | --- | --- | --- | --- | --- | --- | --- | --- | --- | --- |
| | n | A | pA | poly | hom | Ho | He | $\pi$ | $\theta_\pi$ | $\theta_s$ |
| Maui1<br>Nearshore | 18<br>(36) | 31<br>(86%) | 15<br>(83%) | 27<br>(2.2%) | 3<br>(17%) | 0.833 | 0.987 | 0.00596<br>$\pm 0.0032$ | 7.271<br>$\pm 3.87$ | 6.511<br>$\pm 2.25$ |
| Maui1<br>Offshore | 22<br>(44) | 35<br>(80%) | 18<br>(82%) | 31<br>(2.5%) | 4<br>(18%) | 0.818 | 0.986 | 0.00590<br>$\pm 0.0031$ | 7.209<br>$\pm 3.82$ | 7.126<br>$\pm 2.34$ |
| Maui2<br>Nearshore | 23<br>(46) | 38<br>(86%) | 43<br>(93%) | 65<br>(5.3%) | 3<br>(13%) | 0.870 | 0.988 | 0.00614<br>$\pm 0.0035$ | 7.494<br>$\pm 3.96$ | 14.790<br>$\pm 4.48$ |
| Maui2<br>Offshore | 20<br>(40) | 30<br>(75%) | 37<br>(93%) | 33<br>(2.7%) | 4<br>(20%) | 0.800 | 0.967 | 0.00481<br>$\pm 0.0026$ | 5.878<br>$\pm 3.19$ | 7.758<br>$\pm 2.57$ |

#### O‘ahu *vs*. Maui

Inter-island genetic structure, as well as comparison of nearshore and offshore populations were conducted using H2 marker. The hierarchical AMOVA did not detect significant structure between the O‘ahu and Maui populations (*F_CT_*= 0.0069, *P* = 0.272), but the two Maui populations showed significant differentiation (*F_SC_* = 0.0630, *P* = 0.000) based on H2 (Table S2A). The patterns of genetic diversity suggested an overall lower variability in the O‘ahu population; the numbers of haplotypes (A), polymorphic sites (poly), and heterozygous individuals were all smaller on O‘ahu, although nucleotide diversity (π) levels were relatively similar between O‘ahu and Maui (Table 4).

**Table 4.** Population genetic statistics of *P. lobata* from Oʻahu and Maui based on H2 (1221bp). See Table 2 for symbols and abbreviations.

|  | H2 (1221 bp) |  |  |  |  |  |  |  |  |  |
| --- | --- | --- | --- | --- | --- | --- | --- | --- | --- | --- |
| | n | A | pA | poly | hom | Ho | He | $\pi$ | $\theta\pi$ | $\theta s$ |
| Maui1<br>(pooled) | 40<br>(80) | 63<br>(79%) | 76<br>(95%) | 38<br>(3.1%) | 7<br>(15.9%) | 0.825 | 0.986 | 0.0060<br>$\pm 0.0032$ | 7.385<br>$\pm 3.87$ | 7.672<br>$\pm 2.26$ |
| Maui2<br>(pooled) | 43<br>(86) | 68<br>(79%) | 82<br>(95%) | 82<br>(6.7%) | 7<br>(16.2%) | 0.837 | 0.983 | 0.00571<br>$\pm 0.0030$ | 6.972<br>$\pm 3.67$ | 16.316<br>$\pm 4.38$ |
| Maui<br>(pooled) | 83<br>(166) | 126<br>(76%) | 159<br>(81.5%) | 96<br>(7.9%) | 14<br>(16.1%) | 0.831 | 0.987 | 0.00604<br>$\pm 0.0031$ | 7.380<br>$\pm 3.84$ | 16.355<br>$\pm 3.96$ |
| O‘ahu<br>(pooled) | 43<br>(86) | 54<br>(62.7%) | 47<br>(87%) | 35<br>(2.9%) | 9<br>(20.9%) | 0.791 | 0.974 | 0.00643<br>$\pm 0.0033$ | 7.844<br>$\pm 4.08$ | 6.964<br>$\pm 2.06$ |

Pairwise *F_ST_* comparisons between all combinations revealed that the nearshore populations from O‘ahu and Maui1 with a high level of environmental stress were genetically closer to each other than to their respective, nearby offshore populations, and similarly the offshore populations from O‘ahu and Maui1 were genetically closer to each other than to their respective offshore populations (Table 5, O‘ahu and Maui1 populations). Assessing by habitat types, the nearshore and offshore populations of O‘ahu and Maui1 also resulted in significant genetic differentiation (*F_ST_*= 0.0646, *P* = 0.000, Table S2B). The results also revealed that Maui2 corals, which showed no significant structure between the nearshore and offshore sites, turned out to be rather genetically unique compared to the rest of the populations. However, the *F_ST_* values indicated that the Maui2 nearshore population was genetically closer to other offshore populations (*F_ST_*= 0.027-0.035) than to other nearshore populations of O‘ahu and Maui1 (*F_ST_*= 0.139-0.177), suggesting collectively that Maui2 corals at both sites were genetically closer to the offshore populations (Table 5).

**Table 5.** Pairwise *F_ST_* values for all populations from Oʻahu and Maui. The values were estimated using AMOVA in Arlequin with 5000 permutations. Below diagonal = *F_ST_* values, Above diagonal = *P* values. The aster risks refer to the level of statistical significance. N: Nearshore, and O: Offshore.

|  | O‘ahu<br>N | O‘ahu<br>O | Maui1<br>N | Maui1<br>O | Maui2<br>N | Maui2<br>O |
| --- | --- | --- | --- | --- | --- | --- |
| O‘ahu<br>N | - | 0.0002 | 0.3243 | 0.0000 | 0.0000 | 0.0000 |
| O‘ahu<br>O | **0.0791\*\*\*** | - | 0.0079 | 0.6663 | 0.0121 | 0.0462 |
| Maui1<br>N | 0.0006 | **0.0545\*\*** | - | 0.0040 | 0.0000 | 0.0000 |
| Maui1<br>O | **0.0774\*\*\*** | -0.0073 | **0.0517\*\*** | - | 0.0151 | 0.0143 |
| Maui2<br>N | **0.1765\*\*\*** | **0.0348\*** | **0.1387\*\*\*** | **0.0268\*** | - | 0.0716 |
| Maui2<br>O | **0.2047\*\*\*** | **0.0258\*** | **0.1697\*\*\*** | **0.0303\*** | 0.0132 | - |

Genetic structure of *P. lobata* across islands was also visualized using network analysis, which revealed three major clusters of H2 sequences, similar to the results from the Oʻahu populations (Fig. 5). Grouping by habitat-based genetic groups, Cluster 1 was dominated by the offshore type (including Maui2-nearshore) (88 %), Cluster 2 was dominated by nearshore individuals (74%), and Cluster 3 had approximately same proportion of nearshore and offshore types, depicting separation of offshore and nearshore individuals, especially for O‘ahu and Maui1 populations. No clear pattern was observed based on geographic locations (Fig. S3)

**Fig. 5.**
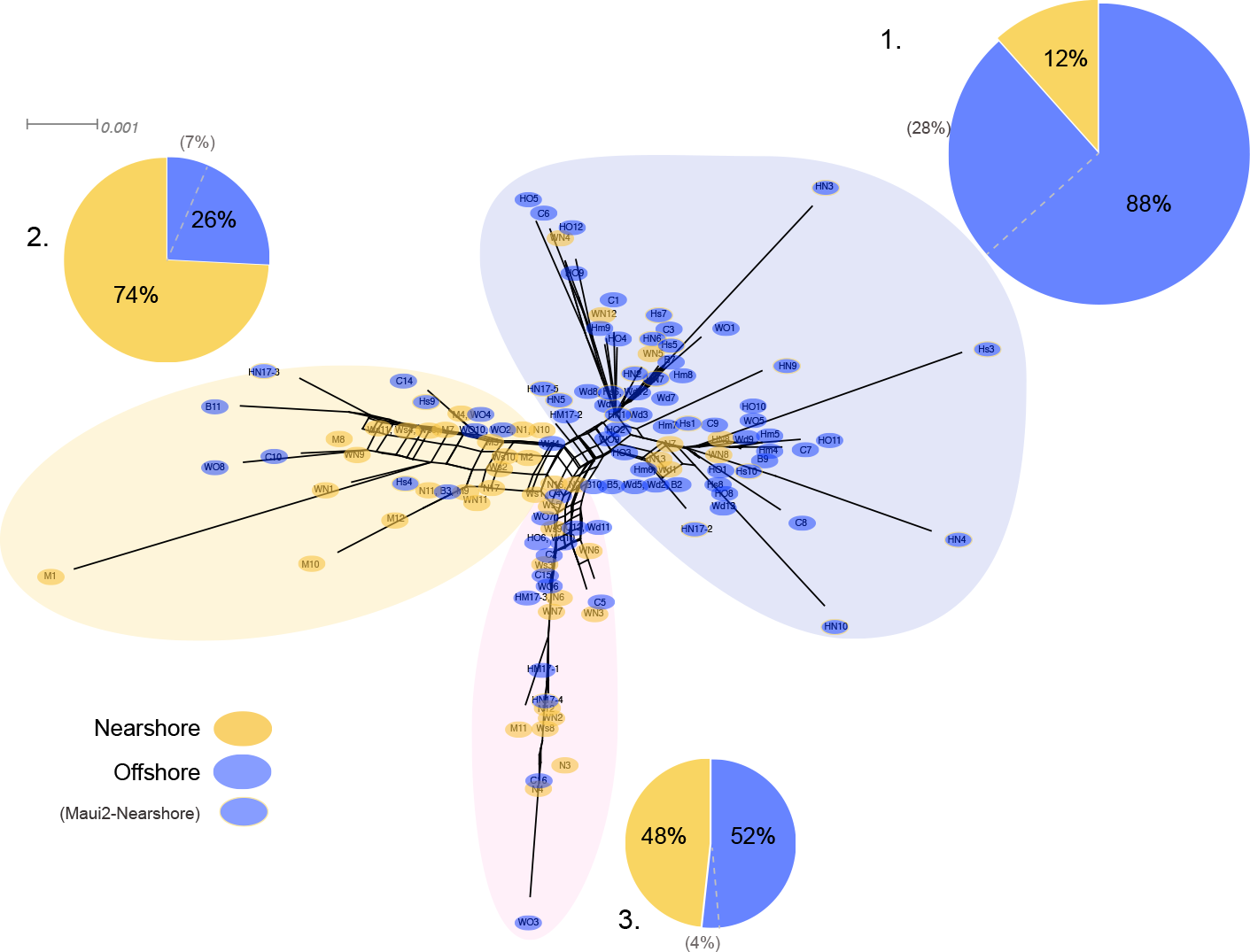
Diagrams of neighbor-net tree networks generated by SplitsTree v.4.14.2 for O‘ahu and Maui *P. lobata* populations based on unphased H2 sequences. Colors are based on habitat-based genetic clusters: Blue color represents the offshore group (including Maui2-nearshore population), and tan color represents the two genetically-close nearshore populations of O’ahu and Maui1. The pie charts show the proportion of sequences present in each group. The gray numbers in () represent the proportion of Maui2 nearshore population.

## Discussion

Previous work at the Archipelagic scale^28^ found IBD, although habitat-level variation was not examined. At a much smaller spatial scale considered here, with explicit sampling of contrasting habitats from across this strong anthropogenic gradient, the pattern of genetic structure we observed for *P. lobata* in Hawaiʻi did not follow IBD (Mantel Test, *r* = −0.0911, *P* = 0.535). Instead, a clear pattern of IBE was revealed with a correlation between habitat types irrespective of geographic distance; the pairwise *F_ST_* values revealed that offshore individuals from two separate islands (>100 km) were genetically closer to each other than to their geographically closest nearshore individuals (300m – 2km), and nearshore individuals from two islands were also typically genetically closer to each other than to adjacent offshore sites (Table 5, Fig. 5).

Similar patterns of genetic structure have been observed in several reef building corals^41,42^. In the case of *Favia fragum*, the “Tall” ecomorph is a seagrass specialist with morphological adaptations to minimize sediment impacts, whereas the “Short” ecomorph shows morphological specializations for coral reef habitats that decrease its fitness in seagrass beds^19^. These traits are highly heritable and divergent selection appears to be driving reproductive isolation among these morphs and ongoing diversification in these incipient species^45^. Here, we find a similar pattern of repeated genetic differentiation across strong ecological gradients driven by anthropogenic impacts (between adjacent sites 300m − 2km apart), but little evidence of differentiation among similar habitats more than 100km distant. Moreover, corals from a control site that lacks such a strong anthropogenically driven environmental gradient (Maui2) did not show the significant genetic structure documented at the other sites, which further supports the likely role of environment in forming the observed genetic patterns (IBE).

It is particularly interesting to find that not only this pattern of clear genetic partitioning within a bay is repeated on a neighbor island, but also the genetic similarity exists between anthropogenically-impacted nearshore populations from the two separate islands that have been exposed to similar environmental pressures (O‘ahu and Maui1). These nearshore sites have experienced deteriorating water and substrate quality due to terrestrial runoff from urbanization of adjacent watersheds over the past century. This environmental decline is likely to limit new recruitment ^46^ and place the population under strong local selection, similar to what was seen in the Caribbean coral *F. fragum*^19,45^. The repeated pattern among bays that share a strong anthropogenic impact gradient also implies that similar selective forces may be operating at both locations; these nearshore coral populations may be selected for their survivorship under local conditions that have been altered by human impacts. Furthermore, there is reduced genetic diversity in the nearshore habitats (Table 2, Table S1), which represents a subset of the standing genetic variation of the larger population, consistent with there being a limited number of individuals capable of surviving in the nearshore habitats.

Additionally, for the O‘ahu populations, differences in microskeletal morphology have been reported between the nearshore and offshore sites^47^. The most noticeable difference was the height of pali (inner vertical skeletal structure that usually exists in a set of eight in *P. lobata*) within a corallite (the structure associated with individual polyps); nearshore corals had taller and more pronounced pali than the offshore ones. The study suggested that the differences might be due to potential beneficial roles played by larger pali in shedding sediments in turbid water^47^, similar to the case of *F. fragum*. Because exact functions and heritability of these traits are unknown, whether the observed morphological differences are due to divergent selection, as in the case of *F. fragum*, cannot be answered at this point. However, correlation between the morphological and genetic distances reported here is consistent with the idea that divergent natural selection is driving such differences.

Additional work is also needed to determine the specific environmental drivers likely to result in selection across these environmental gradients that generate the observed IBE pattern. As discussed earlier, there are many factors, both natural and anthropogenic, that contrast between nearshore and offshore environments (e.g. salinity, irradiance, UV exposure, temperature, pH, wave exposure, nutrients, and biological community) and any of these could contribute to create genetic partitioning between nearshore and offshore sites. For example, a comparable pattern of genetic structure has been observed across a particularly strong temperature gradient between the back-reef and forereef *P. lobata* populations in the areas with negligible terrestrial runoff or pollution in American Samoa^23^. However, in Hawaiʻi we see no such differentiation between nearshore and offshore sites at our unimpacted control (Maui2), supporting the hypothesis that the primary driver of differentiation is anthropogenic. It will be important to continue to observe whether this differentiation is transient and of little evolutionary importance, or whether it progresses towards incipient speciation, as it appears to have done in the Caribbean coral *F. fragum*^19,45^.

Our results show that the broadly distributed broadcast spawning coral *P. lobata* exhibits very fine-scale (300m to 2km) genetic structure, and environmental drivers across habitat types have a stronger effect in forming such genetic structure (IBE) than geographic distances (IBD) in these coastal areas. Without thorough sampling among habitats at small-scales, we could easily overlook such important local genetic differences, and may mistakenly conclude that populations are uniform across the landscape. In fact, this finding may shed light on the common pattern of ‘chaotic genetic patchiness’^48^ so commonly reported among population genetic studies of marine organisms, in which geographically proximate populations show greater genetic differentiation than those from distant sites (e.g.^19,45,47^). Although our samples are from geographically limited locations, our results demonstrate that understanding small-scale genetic variation and diversity provide important information on the ecological basis of genetic diversity and differentiation, which must be understood to effectively implement future coral reef conservation efforts.

The ecological diversification of reef building corals over a small spatial scale, despite ongoing gene flow, also provides a rare example of genetic divergence in the absence of spatial barriers to gene flow, indicating that divergent natural selection can act as an evolutionary driver of reproductive isolation^23,47^. Here, we extend these findings to include *P. lobata* in Hawaii, which shows likely occurrence of similar diversification process across steep environmental gradients driven by anthropogenic impacts. This may represent the initial stages of adaptive diversification, as seen in other marine species from the Hawaiian Archipelago (e.g. limpets^49,50^). There is clearly some genetic connectivity among adjacent islands, congruent to previous studies^28^, and hence the observed divergence across these steep ecological gradients in spite of high dispersal potential^26,51^ appears consistent with the early phases of speciation with gene flow^52^. Whether this initial stage of divergent selection among habitats is transient or has the potential to progress to later stages remains to be seen, but our results and others^49,50^ indicate that this initial stage can be realized even in a broadcasting species with high dispersal potential. Together, these results add to the growing evidence that the initial phase of speciation is possible without geographic isolation, and lend support to the hypothesis that ecological speciation (*sensu*^53^) may be more common in the sea than believed previously.

## Methods

### Species identification

Due to its high morphological plasticity, the genus *Porites* is notorious for its difficulties in distinguishing between its species (e.g.^27, 54–56^). Genetic delineation of some *Porites*, including *P. lobata*, has been challenging due to cryptic species and polymorphic or hybrid species complexes (e.g.^56,57^). Although *Porites* corallites are small, irregular and can be highly variable, micro-skeletal (corallite) structures have been proposed to be more reliable for species identification, therefore, we examined the corallites of all collected samples to confirm our taxonomic identifications^27,57^. In Hawai‘i, the only *Porites* species with a similar colony morphology to *P. lobata* is *Porites evermanni* (there are no records of *Porites lutea* in Hawai‘i, although Fenner^58^ synonymized *P. evermanni* and *P. lutea*, they represent two distinct genetic clades^56^. *P. evermanni* is genetically distinct from *P. lobata*^56^, Clade V), and *P. lutea* has a distinct corallite skeletal morphology, compared to *P. lobata*^27^.

### Coral Sampling

Small fragments (1 cm^2^) of *P. lobata* tissue samples were collected from live colonies between February 2013 to May 2017 at the following sampling sites in Hawaiʻi; a) ‘O‘ahu’-nearshore (n=22) and offshore (n=21) sites at Maunalua Bay, O‘ahu (21.261∼21.278°N, 157.711°W), b) ‘Maui1’ – nearshore (n=21) and offshore (n=23) sites off the Hanakaoʻo Beach Park, West Maui (Wahikuli, 20.95°N, 156.68°W), and c) Maui2 – nearshore (n=23), and offshore (n=20) sites off the Honokōwai Beach Park, West Maui (Honokōwai, 20.90°N, 156.69°W) (Fig. 1, Fig.S1). Samples were taken from coral colonies at least two meters apart at each site, and after sampling, each coral colony was photographed and tagged to avoid resampling of the same colony. The collected tissue samples were either flash frozen in liquid nitrogen on shore and subsequently stored at −80, preserved in DMSO buffer (Gaither et al. 2011), or stored in 100% ethanol. Genomic DNA was extracted from each coral tissue sample using the Qiagen® DNeasy Blood & Tissue Kit. Coral samples were collected under the State of Hawai‘i Division of Aquatic Resources, Special Activity Permit 2013-26, 2014-64, 2015-06, and 2017-16.

### PCR

For the samples from O‘ahu, the following three regions of coral host DNA were PCR-amplified: 1) ∼ 400 bp coral mitochondrial CR with primers CRf and CO3r^59^, 2) ∼ 700 bp coral nuclear ITS with primers ITSZ1 and ITSZ2^56^, and 3) ∼1,500 bp coral nuclear H2 with novel primers zH2AH4f (5’-GTGTACTTGGCTGCYGTRCT-3’) and zH4Fr (5‘- GACAACCGAGAATGTCCGGT-3’). H2 was developed to create a genetic marker that allow direct sequencing of post PCR products to efficiently assess small-scale population genetic structure, because 1) the mitochondrial genome of *P. lobata* exhibits very little sequence variability (< 0.02% polymorphic sites^60^ due to its extremely slow evolutionary rate^44^, and 2) even though high polymorphism in ITS is a desirable trait, sequencing of ITS requires time-consuming cloning, and analyzing the multi-copy gene poses analytical challenges, as it deviates from a standard diploid model. H2 was amplified under the following conditions: 96 °C for 2 min (one cycle), followed by 34 cycles consisting of 96 °C for 20 s, 58.5 °C for 20 s, and 72 °C for 90 s, and a final extension at 72 °C for 5 min. H2 amplifications (25 µl) consisted of 0.5 µl of DNA template, 0.2 µl of GoTaq^®^ DNA Polymerase (Promega, Madison, WI), 5 µl of GoTaq^®^ Reaction Buffer, 1.6 µl of 50mM MgCl_2_, 2 µl of 10 mM dNTPmix, 1.6 µl of each 10mM primer, and nuclease-free water to volume. For samples with multiple bands, approximately 1500-bp PCR products were extracted from agarose gels after electrophoresis and purified using the UltraClean® 15 DNA Purification Kit (MO BIO Laboratories, Carlsbad, CA) according to the manufacturer’s instruction. The rest of the PCR products were purified with UltraClean® PCR Clean-Up Kit (MO BIO Laboratories) and sequenced directly in both directions on the ABI 3730xl DNA Analyzer. Clone libraries were created for each amplified ITS region using the pGEM^®^-Easy Vector System (Promega). Positive inserts were verified by PCR using SP6 and T7 primers, and plasmids (2–5 per library) were treated with UltraClean® 6 Minute Mini Plasmid Prep Kit (MO BIO Laboratories) and sequenced on an ABI-3130XL Genetic Analyzer sequencer. For Maui samples, H2 was amplified and sequenced using the same method as described above.

### Sequence analyses

Resulting DNA sequences were aligned using Geneious^®^ 6.1.8 (Biomatters Ltd., Auckland, New Zealand). Polymorphic sites within H2 regions were identified using Geneious® (Find Heterozygotes option) and confirmed by eye. Middle sections, as well as both ends of H2 were then trimmed to 1,352 bp (for O‘ahu sequences) or 1,221 bp (for combined O‘ahu and Maui analysis) due to many having low quality and/or missing nucleotides. H2 was phased using the program PHASE 2.1 ^61^ and SeqPHASE ^62^. The analysis of molecular variance and other population genetic statistics were estimated in Arlequin 3.5 ^43^ and TCS 1.21 ^63^. The global AMOVA with a weighted average over loci with permutation tests was used as implemented in Arlequin 3.5. For H2, both phased and non-phased sequences were run with AMOVA, which produced the same statistical results, and therefore only the results from the phased sequences are presented here. Up to five coral ITS sequences were successfully cloned and sequenced per colony, and the entire data set was used for calculation of population statistics, treating each cloned sequence as a haplotype. Attempts have been made to conduct genetic analysis using ITS by a) treating each sequence as a haplotype (inclusivity), b) making a consensus sequence per individual (consensus by plurality), or c) using a hierarchal PERMANOVA ^23^. In this study, we ran AMOVA using ITS by both a) and b) methods, which produced the same statistical outcome, and hence, the results from inclusivity (a) are presented in this paper. To address the unequal sample sizes (28 vs 44) between the sites in Maunalua Bay, the analysis was repeated after resampling to the equal sample size (28) for 10 times. All DNA sequences were inspected for possibility of multi-sampled individuals, and all sampled colonies were considered as separate individuals (genets) since no two individuals from a single site shared the same haplotypes (H2). Mantel’s test for isolation by distance was run on the samples in R^64^ using pairwise genetic distance with 5000 bootstrap permutations. Rarefaction anaysis was conducted in Analytic Rarefaction 2.1.1^65^.

## Acknowledgements

We thank the following people for assistance in collecting corals: M. Stiber, V. Sindorf, F. Seneca, J. Murphy, J. Martinez, A. Lyman, K. Richmond, V. Hölzer, N. Spies, and A. Irvine. Special thanks to T. Oliver for their support and advice. ZHF would like to thank the Seaver Institute for support, and ZHT and RJT were also supported by NSF OA#14-16889. This is HIMB contribution #xxxx and SOEST contribution #xxxx.

## Author contributions

KHT designed research, performed research, and analyzed data. ZHF and RJT contributed to designing research and analyzing data. RHR contributed to designing research, funds, analytical tools and reagents. KHT wrote the paper with major contributions, suggestions, and approval from all authors.

## Data Availability

All DNA sequences are available from GenBank (ITS: Accession# KY493091- KY493160, H2: Accession# KY502280 - KY 502376 and MF629151-MF629152, CR: Accession# KY502373- KY502375). The phased H2 sequences used in the analysis are included as S1 File. All other data are available upon request.

## Supplementary Information

**Fig. S1.**
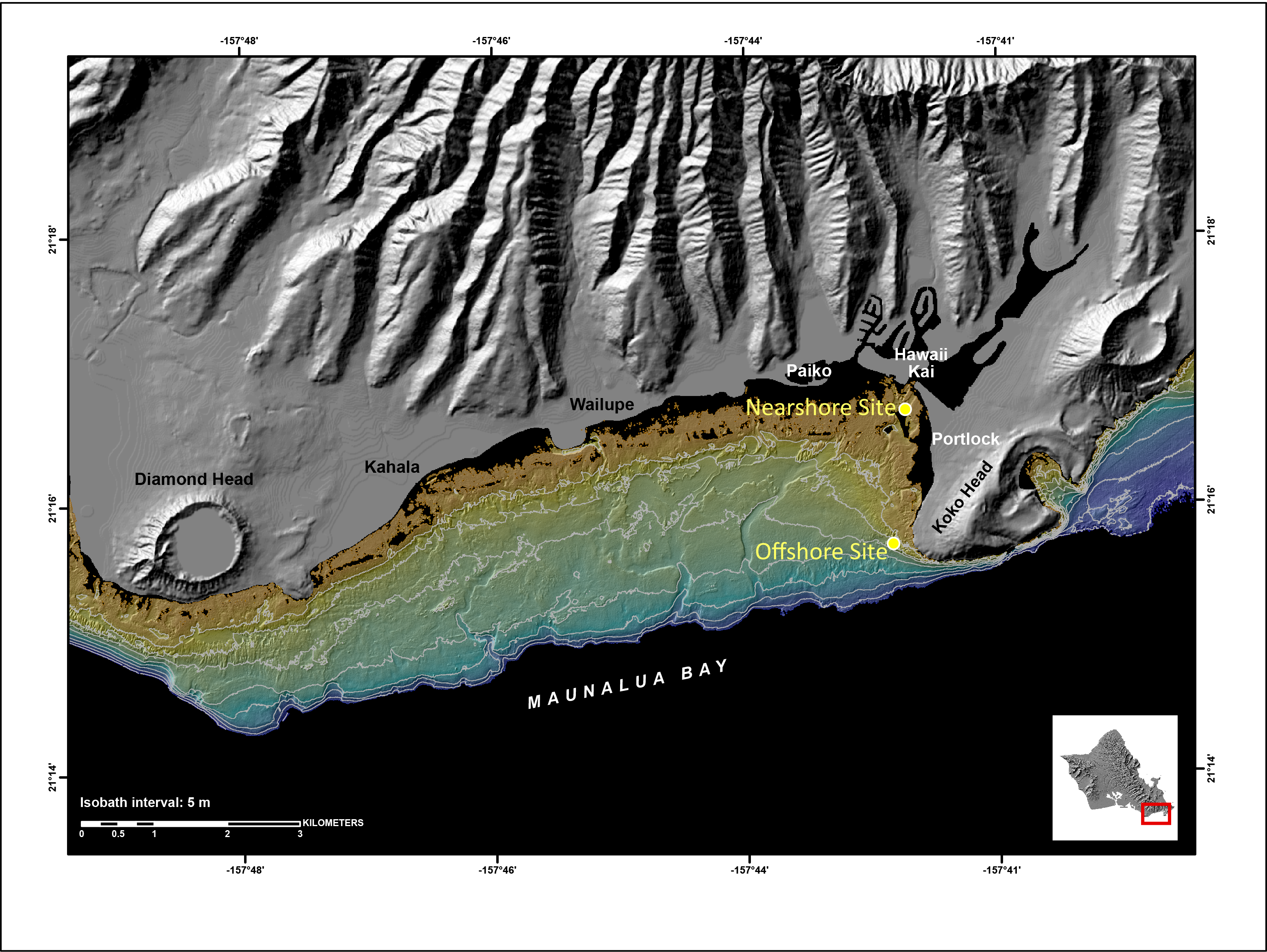
Map of Maunalua Bay with bathymetry, showing the nearshore and offshore sampling locations (Map was provided by Curt Storlazzi, U.S. Geological Survey).

**Fig. S2.**
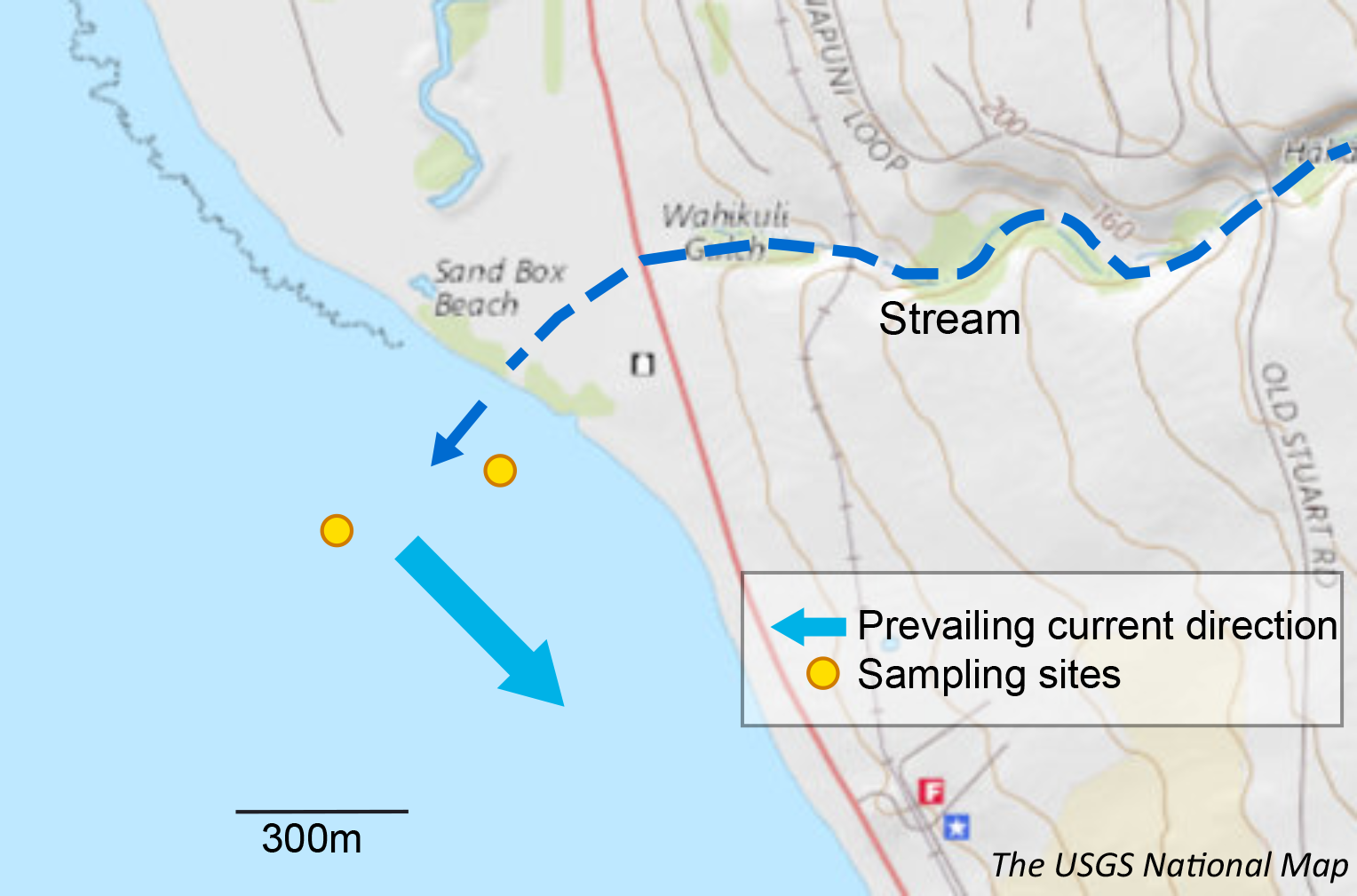
Map of Maui1 (Wahikuli) sampling location, showing that the run-off point that drains turbid water, and the unique current patterns that create clear differences in water quality (Map services and data available from U.S. Geological Survey, National Geospatial Program).

**Fig. S3.**
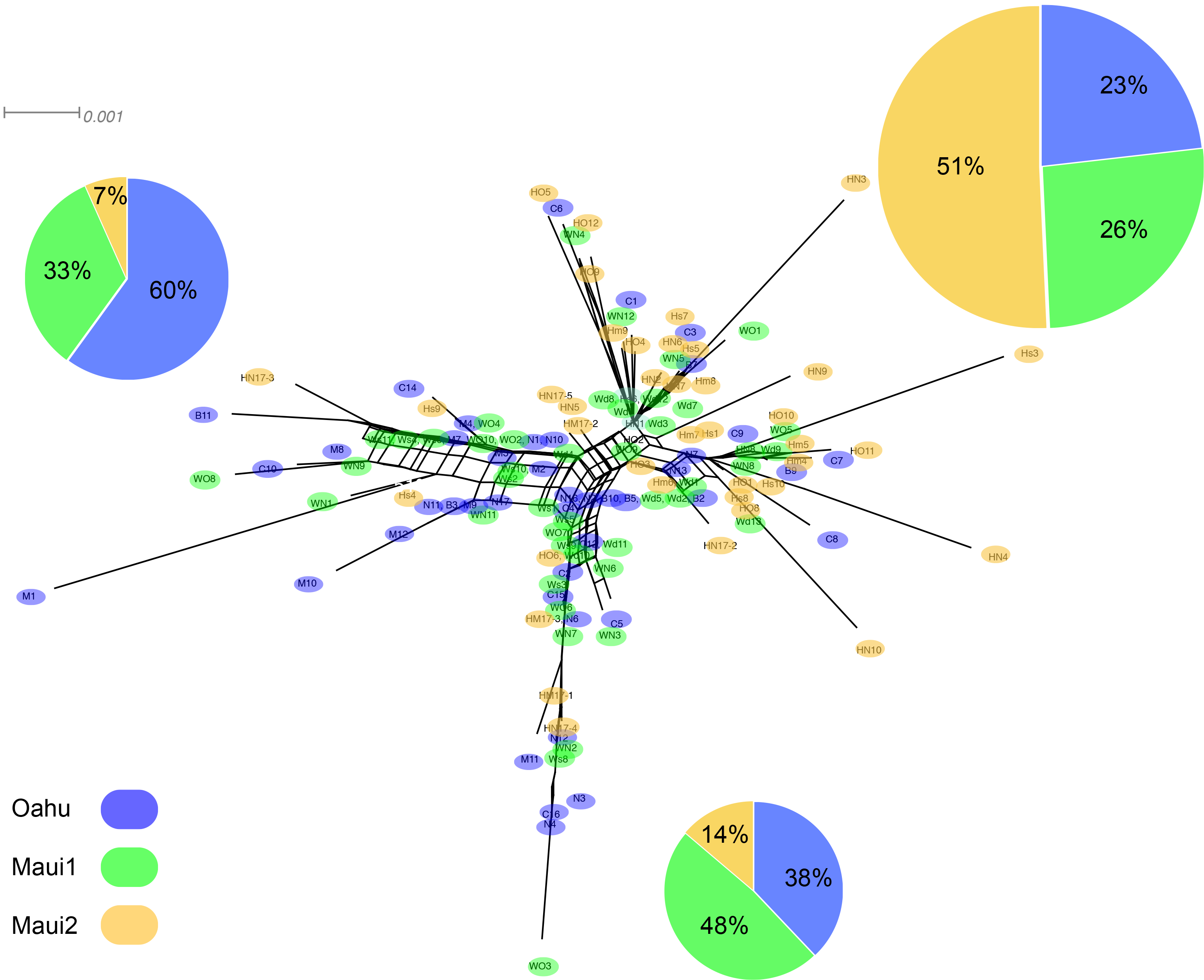
Diagrams of neighbor-net tree networks generated by SplitsTree v.4.14.2 for O‘ahu and Maui *P. lobata* populations based on unphased H2 sequences. Colors are based on geographic locations.

**Table S1.**
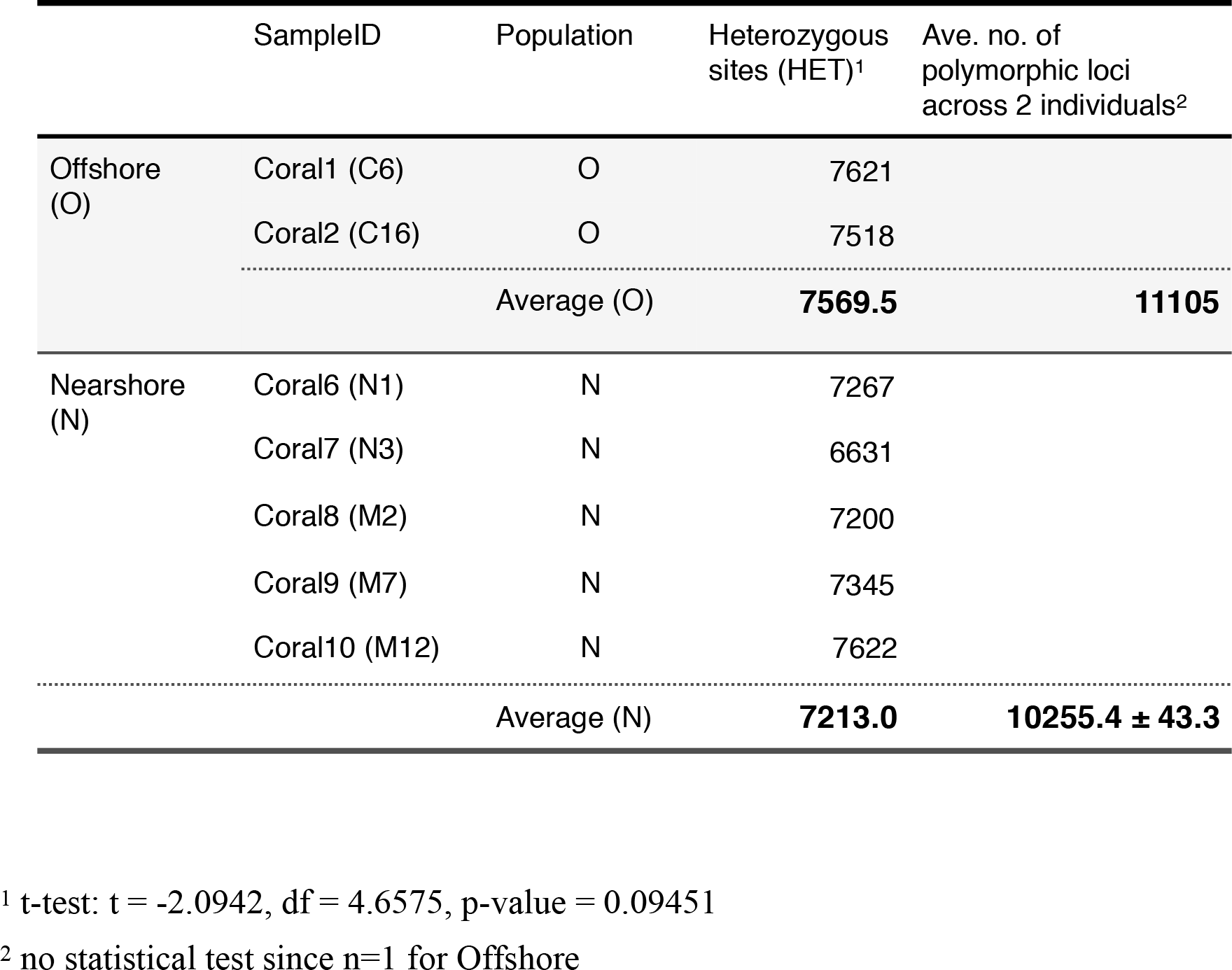
Genetic diversity comparison between the nearshore and offshore *P. lobata* populations from Maunalua Bay, Hawaii (Oʻahu) based on 17,850 single nucleotide polymorphic loci. The number of heterozygous sites per individual was obtained using VCFtools, and the total number of polymorphic loci was obtained using Arlequin (for the nearshore population, showing the average over two individuals).

**Table S2.** AMOVA results of *P. lobata* populations across islands (A), and across habitat types: nearshore vs offshore pooled individuals from Oʻahu and Maui1 (B), and Oʻahu and Maui 1 & 2 (C).

| A | Source of Variation | Variance components | % Variance | Fixation indices |
| --- | --- | --- | --- | --- |
| | Among groups (islands) | 0.00135 | 0.03 | $F_{CT} = 0.00034$ |
| | Among populations within groups | 0.24897 | 6.35 | $F_{SC} = \mathbf{0.06349^{***}}$ |
| | Among individuals Within populations | 1.51485 | 38.62 | $F_{IS} = \mathbf{0.41248^{***}}$ |
| | Within individuals | 2.15769 | 55.00 | $F_{IT} = \mathbf{0.44997^{***}}$ |
| B | Source of Variation | Variance components | % Variance | $F_{ST}$ |
| | Between sites | 0.2360 | 5.96 | $\mathbf{0.0596^{***}}$ |
|  | Within sites | 1.3906 | 35.12 |  |
|  | Within individuals | 2.3333 | 58.92 |  |
| C | Source of Variation | Variance components | % Variance | $F_{ST}$ |
| | Between sites | 0.1074 | 2.73 | $\mathbf{0.0273^{***}}$ |
|  | Within sites | 1.6703 | 42.44 |  |
|  | Within individuals | 2.1577 | 54.82 |  |

**S1 File. Supporting information.** Phased H2 sequences of all individuals used in the analysis.

## References

1. Hughes, T. P., Graham, N. A. J., Jackson, J. B. C., Mumby, P. J. & Steneck, R. S. Rising to the challenge of sustaining coral reef resilience. TRENDS in Ecology and Evolution 25, 633–642 (2010).

2. Graham, N. A. J. Habitat Complexity: Coral Structural Loss Leads to Fisheries Declines. Current Biology 24, R359–R361 (2014).

3. Richmond, R. H. & Wolanski, E. in Corals Reefs an Ecosystem in Transition (eds. Dubinsky, Z. & Stambler, N.) 3–12 (Springer, Dordrecht, 2011).

4. Wenger, A. S. et al. Effects of reduced water quality on coral reefs in and out of no-take marine reserves. Conservation Biology 30, 142–153 (2015).

5. Smith, T. B. et al. Assessing coral reef health across onshore to offshore stress gradients in the US Virgin Islands. MPB 56, 1983–1991 (2008).

6. van Dam, J. W., Negri, A. P., Uthicke, S. & Mueller, J. F. Chemical pollution on coral reefs: exposure and ecological effects, in Ecological impact of toxic chemicals (eds. Sánchez-Bayo, F., van den Brink, P. J. & Mann, R. M.) 187–211 (Bentham Science Publishers Ltd., 2011).

7. Thompson, A., Schroeder, T., Brando, V. E. & Schaffelke, B. Coral community responses to declining water quality: Whitsunday Islands, Great Barrier Reef, Australia. Coral Reefs 33, 923–938 (2014).

8. Ennis, R. S., Brandt, M. E., Grimes, K. R. W. & Smith, T. B. Coral reef health response to chronic and acute changes in water quality in St. Thomas, United States Virgin Islands. Marine Pollution Bulletin 111, 418–427 (2016).

9. Gorospe, K. D. & Karl, S. A. Small-scale spatial analysis of in situ sea temperature throughout a single coral patch reef. Journal of Marine Biology 2011, 1–12 (2011).

10. Guadayol, Ò., Silbiger, N. J., Donahue, M. J. & Thomas, F. I. M. Patterns in temporal variability of temperature, oxygen and pH along an environmental gradient in a coral reef. PLoS ONE 9, e85213 (2014).

11. Morgan, K. M., Perry, C. T., Smithers, S. G., Johnson, J. A. & Daniell, J. J. Evidence of extensive reef development and high coral cover in nearshore environments: implications for understanding coral adaptation in turbid settings. Sci. Rep. 6, 29616 (2016).

12. Wang, I. J. & Bradburd, G. S. Isolation by environment. Molecular Ecology 23, 5649–5662 (2014).

13. Wright, S. Isolation by distance. Genetics 28, 114–138 (1943).

14. Selkoe, K. A. et al. A decade of seascape genetics: contributions to basic and applied marine connectivity. Mar. Ecol. Prog. Ser. 554, 1–19 (2016).

15. Orsini, L., Vanoverbeke, J., Swillen, I., Mergeay, J. & De Meester, L. Drivers of population genetic differentiation in the wild: isolation by dispersal limitation, isolation by adaptation and isolation by colonization. Molecular Ecology 22, 5983–5999 (2013).

16. Nosil, P., Funk, D. J. & ORTIZ-BARRIENTOS, D. Divergent selection and heterogeneous genomic divergence. Molecular Ecology 18, 375–402 (2009).

17. De Meester, L., Gómez, A., Okamura, B. & Schwenk, K. The Monopolization Hypothesis and the dispersal–gene flow paradox in aquatic organisms. Acta oecologica 23, 121–135 (2002).

18. McRae, B. H. & Beier, P. Circuit theory predicts gene flow in plant and animal populations. Proc. Natl. Acad. Sci. U.S.A. 104, 19885–19890 (2007).

19. Carlon, D. B. & Budd, A. F. Incipient speciation across a depth gradient in a scleractinian coral? Evolution 56, 2227–2242 (2002).

20. Carlon, D. B., Budd, A. F., Lippé, C. & Andrew, R. L. The quantitative genetics of incipient speciation: heritability and genetic correlations of skeletal traits in populations of diverging *Favia fragum* ecomorphs. Evolution 65, 3428–3447 (2011).

21. Rundle, H. D. & Nosil, P. Ecological speciation. Evolution 8, 336–352 (2005).

22. Bongaerts, P. et al. Genetic Divergence across Habitats in the Widespread Coral *Seriatopora hystrix* and Its Associated *Symbiodinium*. PLoS ONE 5, e10871 (2010).

23. Barshis, D. J. et al. Protein expression and genetic structure of the coral *Porites lobata* in an environmentally extreme Samoan back reef: does host genotype limit phenotypic plasticity? Molecular Ecology 19, 140297–140297 (2010).

24. Wolanski, E., Martinez, J. A. & Richmond, R. H. Quantifying the impact of watershed urbanization on a coral reef: Maunalua Bay, Hawaii. Estuarine, Coastal and Shelf Science 84, 259–268 (2009).

25. Richmond, R. H. HCRI Project Report (FY2009-2010): Watersheds impacts on coral reefs in Maunalua Bay, Oahu, Hawaii. 1–8 (Hawaii Coral Reef Initiatives, 2011).

26. Presto, K. M., Storlazzi, C. D., Logan, J. B., Reiss, T. E. & Rosenberger, K. J. Coastal Circulation and Potential Coral-larval Dispersal in Maunalua Bay, Oahu, Hawaii - Measurements of waves, Currents, Temperature, and salinity June-September 2010. 1–67 (U.S. Geological Survey Open-File Report 2012-1040, 2012).

27. Veron, J. E. N. Corals of the World. 1–3, (Australian Institute of Marine Science, 2000).

28. Polato, N. R., Concepcion, G. T., Toonen, R. J. & Baums, I. B. Isolation by distance across the Hawaiian Archipelago in the reef-building coral Porites lobata. Molecular Ecology 19, 4661–4677 (2010).

29. Baums, I. B., Boulay, J. N., Polato, N. R. & Hellberg, M. E. No gene flow across the Eastern Pacific Barrier in the reef-building coral *Porites lobata*. Molecular Ecology 21, 5418–5433 (2012).

30. Stafford-Smith, M. G. Sediment-rejection efficiency of 22 species of Australian scleractinian corals. Marine Biology 115, 229–243 (1993).

31. Levas, S. J., Grottoli, A. G., Hughes, A., Osburn, C. L. & Matsui, Y. Physiological and Biogeochemical Traits of Bleaching and Recovery in the Mounding Species of Coral Porites lobata: Implications for Resilience in Mounding Corals. PLoS ONE 8, e63267 (2013).

32. Roff, G. et al. Porites and the Phoenix effect: unprecedented recovery after a mass coral bleaching event at Rangiroa Atoll, French Polynesia. Marine Biology 161, 1385–1393 (2014).

33. Franklin, E. C., Jokiel, P. L. & Donahue, M. J. Predictive modeling of coral distribution and abundance in the Hawaiian Islands. Marine ecology Progress Series 481, 121–132 (2013).

34. LaJeunesse, T. et al. High diversity and host specificity observed among symbiotic dinoflagellates in reef coral communities from Hawaii. Coral Reefs 23, 596–603 (2004).

35. Smith, L. W., Wirshing, H. H., Baker, A. C. & Birkeland, C. Environmental versus genetic influences on growth rates of the corals Pocillopora eydouxi and Porites lobata (Anthozoa: Scleractinia) 1. Pacific Science 62, 57–69 (2008).

36. Fabina, N. S., Putnam, H. M., Franklin, E. C., Stat, M. & Gates, R. D. Transmission mode predicts specificity and interaction patterns in coral-*Symbiodinium* networks. PLoS ONE 7, e44970 (2012).

37. Richmond, R. H. HCRI Project Progress Report FY2008: Watersheds impacts on coral reefs in Maunalua Bay, Oahu, Hawaii. 1–9 (2009).

38. Storlazzi, C. D., Presto, K. M., Logan, J. B. & Field, M. E. Coastal Circulation and Sediment Dynamics in Maunalua Bay, Oahu, Hawaii. 1–64 (USGS Open-File Report 2010-1217, 2010).

39. Rodgers, K. S., Jokiel, P. L., Brown, E. K., Hau, S. & Sparks, R. Over a Decade of Change in Spatial and Temporal Dynamics of Hawaiian Coral Reef Communities 1. Pacific Science 69, 1–13 (2015).

40. Williams, I., Vargas, B., White, D. & Callender, T. West Maui Wahikuli & Honokowai Priority Watershed Area: Reef Condition Report. 1–18 (Sceintific Report by Hawaii Division of Aquatic Resources, Ridge 2 Reef Initiative, NOAA Coral Reef Conservation Program., 2014).

41. Storlazzi, C. D., McManus, M. A., Logan, J. B. & McLaughlin, B. E. Cross-shore velocity shear, eddies and heterogeneity in water column properties over fringing coral reefs: West Maui, Hawaii. Continental Shelf Research 26, 401–421 (2006).

42. Vargas-Angel, B. Maui_Baseline_report_2017. 1–49 (2017). doi:10.7289/V5/SP-PIFSC-17-001

43. Excoffier, L. & Lischer, H. E. L. Arlequin suite ver 3.5: a new series of programs to perform population genetics analyses under Linux and Windows. Molecular Ecology Resources 10, 564–567 (2010).

44. Shearer, T. L., van Oppen, M. J. H., Romano, S. L. & Wörheide, G. Slow mitochondrial DNA sequence evolution in the Anthozoa (Cnidaria). Molecular Ecology 11, 2475–2487 (2002).

45. Carlon, D. B. & Lippé, C. Estimation of mating systems in Short and Tall ecomorphs of the coral Favia fragum. Molecular Ecology 20, 812–828 (2011).

46. Puritz, J. B. & Toonen, R. J. Coastal pollution limits pelagic larval dispersal. Nature Commuunications 2, 226 (2011).

47. Tisthammer, K. H. & Richmond, R. H. Corallite skeletal morphological variation in Hawaiian Porites lobata. Coral Reefs 65, 560–14 (2018).

48. Johnson, M. S. & Black, R. Chaotic genetic patchiness in an intertidal limpet, Siphonaria sp. Marine Biology 70, 157–164 (1982).

49. Bird, C. E. Morphological and Behavioral Evidence for Adaptive Diversification of Sympatric Hawaiian Limpets (Cellana spp.). Integrative and Comparative Biology 51, 466–473 (2011).

50. Bird, C. E., Holland, B. S., bowen, B. W. & Toonen, R. J. Diversification of sympatric broadcast-spawning limpets (Cellana spp.) within the Hawaiian archipelago. Molecular Ecology 20, 2128–2141 (2011).

51. Storlazzi, C. D. & Field, M. E. Winds, waves, tides, and the resulting flow patterns and fuxes of water, sediment, and coral larvae off West Maui, Hawaii. 1–13 (USGS Open-File Report 2008-1215, 2008).

52. Feder, J. L., Egan, S. P. & Nosil, P. The genomics of speciation-with-gene-flow. Trends in Genetics 28, 342–350 (2012).

53. bowen, B. W., Rocha, L. A., Toonen, R. J., Karl, S. A. & Laboratory, T. T. The origins of tropical marine biodiversity. TRENDS in Ecology and Evolution 28, 359–366 (2013).

54. Veron, J. E. N. Corals in Space and Time: The Biogeography and Evolution of the Scleractinia. (Cornell University Press, 1995).

55. Forsman, Z., Wellington, G. M., Fox, G. E. & Toonen, R. J. Clues to unraveling the coral species problem: distinguishing species from geographic variation in Poritesacross the Pacific with molecular markers and microskeletal traits. PeerJ 3, e751 (2015).

56. Forsman, Z. H., Barshis, D. J., Hunter, C. L. & Toonen, R. J. Shape-shifting corals: molecular markers show morphology is evolutionarily plastic in Porites. BMC Evolutionary Biology 9, 45 (2009).

57. Veron, J. E. N. & Pichon, M. Scleractinia of Eastern Australia Part IV Family Poritidae. 1–168 (Australian National University Press, 1982).

58. Fenner, D. Corals of Hawaii. (Mutual Publishing, LLC, 2005).

59. Vollmer, S. V. & Palumbi, S. R. Hybridization and the evolution of reef coral diversity. Science 296, 2023–2025 (2002).

60. Tisthammer, K. H. et al. The complete mitochondrial genome of the lobe coral Porites lobata(Anthozoa: Scleractinia) sequenced using ezRAD. Mitochondrial DNA Part B 1, 247–249 (2016).

61. Stephens, M., Smith, N. J. & Donnelly, P. A new statistical method for haplotype reconstruction from population data. Am. J. Hum. Genet. 68, 978–989 (2001).

62. Flot, J. F. SeqPHASE: a web tool for interconverting PHASE input/output files and FASTA sequence alignments. Molecular Ecology Resources 10, 162–166 (2010).

63. Clement, M., Posada, D. & Crandall, K. A. TCS: a computer program to estimate gene genealogies. Molecular Ecology 9, 1657–1659 (2000).

64. R Core Team. R: A Language and Environment for StatisticalComputing. (R Foundation for Statistical Computing, 2014).

65. Holland, S. Analytic Rarefaction, Version 2, software, Hunt Mountain Software. (2009).

